# Modeling the Impact of Lock-down on COVID-19 Spread in Malaysia

**DOI:** 10.1101/2020.07.17.208371

**Authors:** Altahir A. Altahir, Nirbhay Mathur, Loshini Thiruchelvam, Ghulam E. Mustafa Abro, Syaimaa’ S. M. Radzi, Sarat C. Dass, Balvinder Singh Gill, P. Sebastian, Saiful A.M. Zulkifli, Vijanth S. Asirvadam

## Abstract

After a breakdown notified in Wuhan, China in December 2019, COVID-19 is declared as pandemic diseases. To the date more than 13 million confirmed cases and more than half a million are dead around the world. This virus also attached Malaysia in its immature stage where 8718 cases were confirmed and 122 were declared as death. Malaysia responsibly controlled the spread by enforcing MCO. Hence, it is required to visualize the pattern of Covid-19 spread. Also, it is necessary to estimate the impact of the enforced prevention measures. In this paper, an infectious disease dynamic modeling (SEIR) is used to estimate the epidemic spread in Malaysia. The main assumption is to update the reproduction number Rt with respect to the implemented prevention measures. For a time-frame of five month, the Rt was assumed to vary between 2.9 and 0.3. Moreover, the manuscript includes two possible scenarios: the first will be the extension of the stricter measures all over the country, and the second will be the gradual lift of the lock-down. After implementing several stages of lock-down we have found that the estimated values of the *Rt* with respect to the strictness degree varies between 0.2 to 1.1. A continuous strict lock-down may reduce the *Rt* to 0.2 and accordingly the estimated active cases will be reduced to 20 by the beginning of September 2020. In contrast, the second scenario considers a gradual lift of the enforced prevention measures by the end of June 2020, here we have considered three possible outcomes according to the MCO relaxation. Thus, the estimated values of *Rt* = 0.7, 0.9, 1.1, which shows a rapid increase in the number of active cases. The implemented SEIR model shows a close resemblance with the actual data recorded from 10, March till 7, July 2020.

**Author summary:** Conceptualization, A.A.A; methodology, A.A.A, N.M; validation, A.A.A, N.M; formal analysis, A.A.A; investigation, N.M, A.A.A; resources, G.E.M.A, L.T; data collection, L.T, N.M; writing—original draft preparation, A.A.A, L.T, G.E.M.A, N.M; writing—review and editing, V.S.A, S.C.D, B.S.G, P.S, S.A.B.M.Z, N.M; visualization, N.M; supervision, V.S.A; project administration, V.S.A. All authors have read and agreed to the published version of the manuscript

## Introduction

COVID-19 is one of the coronaviruses, originated from a large family of viruses, causing illness to animals and humans. These corona viruses trigger a respiratory infection ranging from basic cold to more severe diseases, such as Middle East Respiratory Syndrome (MERS) and Severe Acute Respiratory Syndrome (SARS). The strain of the virus underlying this COVID-19 illness is known as the Severe Acute Respiratory Syndrome corona virus 2 (SARS-CoV-2). The current strain of the virus is first discovered in Wuhan, China, in December 2019 and further spread to other countries [1–3].

Common symptoms of COVID-19 are cough, fever, sore throat, pneumonia, and shortness of breath [4–6]. All groups of people can be affected by this illness; however, about half of them needs a hospital admission due to medical problem backgrounds such as high blood pressure, heart and diabetes [7–9].

The COVID-19 disease spreads from an infected person to an uninfected person through breathing small droplets from nose or mouth, expelled when the COVID-19 infected person coughs, sneezes, or speaks. These droplets do not sink to the ground nor travel far. Thus, nearby people can also get infected when they first touch the surfaces with virus droplets and then touch their face, eyes, nose, or mouth. Therefore, it is advised to always wash hand too [7, 10, 11].

The chronology of COVID-19 spread starts when China’s Health Authority reported a mysterious pneumonia found among 41patients from Wuhan city, on 31st December 2019. This first cluster is linked to the Huanan Seafood Wholesale Market [5, 12, 13]. Accordingly, Wuhan city is placed under quarantine and followed by the whole province of Hubei in the next days [4, 13]. However, few days later China recorded its first case of death [14]. The coronavirus case and even case of death also starts to be observable in regions outside of China. Other than China, COVID-19 was also found to start having outbreak in other countries, South Korea, Iran, Italy, and Spain. On March 11, the WHO has declared COVID-19 as a pandemic [11, 14–16].

As for Malaysia, the Malaysian authorities have initiated a thermal checkup at all the entry points of the country since the reporting of the first wave of the virus. The very first recorded case COVID-19 is linked to a male returning from Wuhan, on the 25^*th*^ January 2020 [17]. Following that, on 27^*th*^ January, all the travelers from Hubei province are banned from entering the country. On 9^*th*^ February, the ban is extended to include all travelers from Jiangsu and Zhejiang provinces [49] and on 5th March, Malaysia added seven regions towards its travel ban order. The travel ban is extended to include Italy, Iran, and South Korea on 11th March, while Malaysians coming from those countries will be quarantined for 14 days. On 12th of March, the Malaysian Authorities announced the first case of human to human transmission, occurring within the local population. As a response to the increasing number of cases, the Malaysian authorities announced the Movement Control Order (MCO) on 16 March 2020. Accordingly, six restrictions have been imposed [50].

1. Mass gathering, and massive events are prohibited (i.e. religious, sports, social and cultural activities).
2. A health screening and self-quarantine for 14 days are imposed on Malaysians returning from abroad.
3. Tourists and foreign visitors are restricted to enter the country.
4. Closure of all kindergartens, government, and private schools.
5. Closure of all public and private higher education institution (IPTs) and skill training institutes.
6. Closure of all government and private premises except for essential services.

With the movement control order put in place since 18^*th*^ March, all citizens have been prohibited from leaving the country and foreigners also prohibited from entering the country. The movement control order was originally set to end on 31st March; however, the order has been extended three times as additional two-week “stage”over the course of two months. For the purpose of estimating the active cases in Malaysia, the various stages are described as follows:

- Stage 0: this stage considers the time period started on the major outbreak occurred on 10 March 2020 till the implementation of the MCO on 18 March 2020.since the discovery of the first case.
- Stage 1: the implementation of the movement control order (MCO) which started on 18 March and set to end on 31 March 2020.
- Stage 2: as new cases continue to raise, the extension of the MCO is announced on 25 March to 14 April. This stage is known as Enhanced Movement Control Order (EMCO), where specific locations were subjected to a stricter order. Accordingly, all residents and visitors within those areas are forbidden from exiting their homes during the order; non-residents and visitors outside the area cannot enter into those areas; all businesses are shut down and all roads into the areas are blocked; and adequate food supplies were given by the authorities during the 14 day-order to all residents.
- Stage 3: according to the WHO projection of the peak which estimated to take place in mid-April, the third extension is announced on 10 April, which extends the MCO to 28 April.
- Stage 4: announced on 23 April and extends the MCO to 12 May.
- Stage 5: This stage considers the time period between 12 - 30 May 2020. Accordingly, the government announced a plan known as Conditional Movement Control Order (CMCO), aims to relax the enforced regulations in order to reopen the national economy in a controlled manner [44]. The regulations of the CMCO were eased where most economic sectors and activities are allowed to operate while observing the business standard operation. However, sports activities involving large gatherings, social, community, cultural and religious events, as well as all types of official events and assemblies are not permitted [44, 45].
- Stage 6: On 7 June, the Malaysian Authorities announced the Recovery Movement Control Order (RMCO) between 10 June and 31 August. Accordingly, the government permits the interstate travel except for the areas remaining under the Enhanced Movement Control Order (EMCO). In most areas, certain religious activities at mosques are allowed again, but with additional restrictions. On 26 June, the Malaysian Authorities announced more relaxation regarding cultural and tourism related sectors. However, tourism businesses are required to abide the social distancing measures, wear face masks, temperature checkup as well as a crowd limit. On 29 June, it was reported that both government and private pre-schools, kindergartens, nurseries, and day care centers would resume operations from 1 July.

This research article is divided into the following sections: Section 2 (Background), gives overview of COVID-19 in other countries and cities. It also provides a clear description of COVID-19 prediction modeling which has been used by many other country experts to predict values for controlling cases. Section 3 presents the actual data in order to see the pattern of virus spread based on country (Malaysia) data and state data. Section 4 describes the model used to predict the future aspects of virus spread. Section 5 discusses the findings of the research. Finally, section 6 concludes the discussion and demonstrates the recommendations of the manuscript.

## Background

It is believed that any system that has intrinsic number of uncertainties, measurement and time delays, such systems are very complex and difficult to predict. Currently the most contagious corona virus has all such factors within itself and it has swallowed already more than 284,000 lives. In the last few months, researchers from all around the globe has suggested numerous models for predicting the behavior of COVID-19 curve and the people are somehow familiar with such epidemiological system models. The extended calculations in such models are predicting expectations about what the future holds and estimating the situation of a specific jurisdiction but unfortunately, these models have some imperfects within. Discussing the agent-based models, they are depending over the mobility and transportation which allows people to move between two points. In most of the cases, this is generated using the dataset from census and from hospitals. Whereas the compartmental model is based on ordinary differential equations used in several cases such that [22–24].

This is observed while doing literature survey that people have tried to predict the *Ro* by any technical strategy hence in the same regard this paper presents the major modeling schemes that are tried to predict the next upcoming months for specific jurisdiction such that from 06 January 2020 till 11 May 2020. For this time interval proposed manuscript identifies the top 10 frequent studies that may estimate the reproductive co-efficient for this novel coronavirus (COVID-19) from all over the world. It has been observed that very few models are available that focuses specific cities of a country [23] [25] [**?**], whereas others are focused on entire country. In Table 1, it has been seen that mostly the approaches are proposed for entire country and very few are proposed for cities.

**Table 1.** Modeling Approaches for COVID-19 Country and City Wise.

| S.No | Approach | Country | City/Region |
| --- | --- | --- | --- |
| 1 | Stochastic Markov Chain Monte Carlo method [22] | China | – |
| 2 | Mathematical Dynamic Model based on five compartment (SAISTR) [23] | China | – |
| 3 | Statistical Approach [50] | China | Beijing |
| 4 | Mathematical Transmission Model [24] | China | – |
| 5 | Mathematical Exponential Model on serial intervals [25] | China | Wuhan |
| 6 | Mathematical Decay and Exponential Model [27, 28] | China | – |
| 7 | SEIR Modeling with Regression Technique [31–34] | India, South Korea | – |
| 8 | SEIRD Modeling with Metheuristic Technique [35, 36] | South Korea | – |

It has been observed that very few research contributions were made to derive the mathematical model universally applicable to individual countries or locality [24]. Some of the techniques were based on Susceptible, exposed, infectious and recovered SEIR approach [26] and some are based on exponential growth with an amalgamation of statistical approach and stochastic representation [26].

Few estimation techniques are based on the epidemiological parameters which are derived using an age-structured, location and model that uses data on specific contact patterns [29]. Taking an example of Italy which had been affected by COVID-19 recently, people implemented the SEIR Model to compute the future response [37] by varying the parameters and initial conditions. The same model has been adopted by one of the researchers for computing the causalities in the United Kingdom [31]. SEIR Model has a great and efficient computational result, therefore, it has been used by many researchers such that India [30], the data record most of the cases are collected from John Hopkin’s repository in the multiple time frames. In addition to this, several research contributions were made in [31–34] using this SEIR model, therefore, the paper also proposes the same model for computing the casualties in Malaysia.

## Data Visualization of Malaysia for Covid-19

This section will give data representation for the COVID-19 cases that occurred in Malaysia.

### Area of Study

This research only focus for the Coivid-19 analysis in Malaysia. Malaysia is a Southeast Asian country occupying parts of the Malay Peninsula and the island of Borneo. Malaysia has 13 states in total. The 13 states are based on historical Malay kingdoms and 9 of the the 11 Peninsular state known as the Malay states. As shown in Figure 1, the Malaysia map is sketched to show the corona virus spread. In Figure 1 red circles the the hot-spot region where Covid-19 cases are reported.

**Fig 1.**
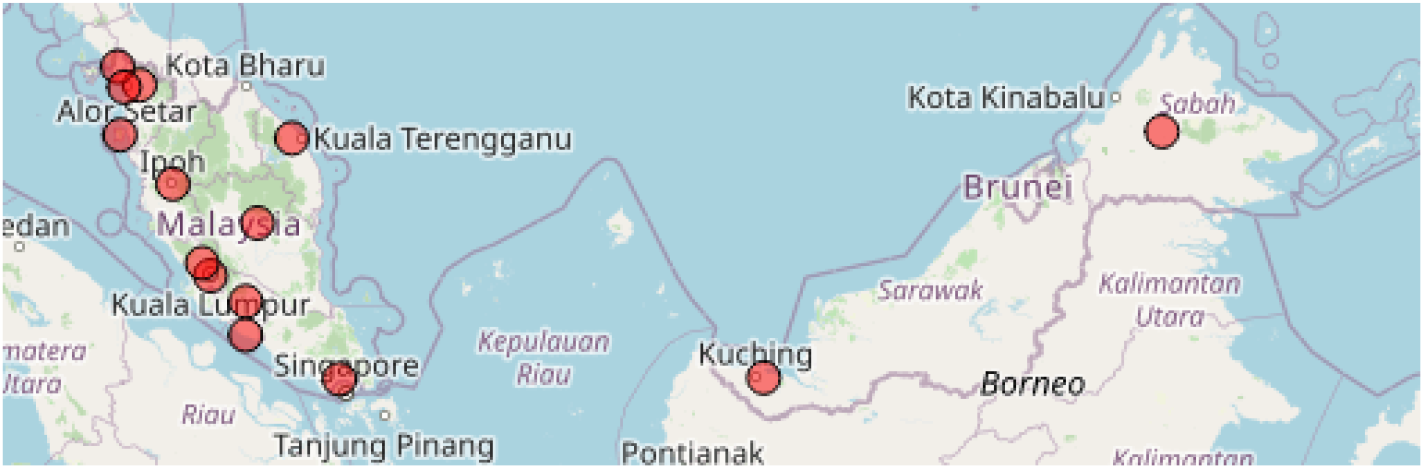
Malaysia map with highlighted cases in red circles.

Figure 2 visualizes the total number of cases count based on 4 categories i.e, Active cases, Cases per day, Total cases, and Total death.

**Fig 2.**
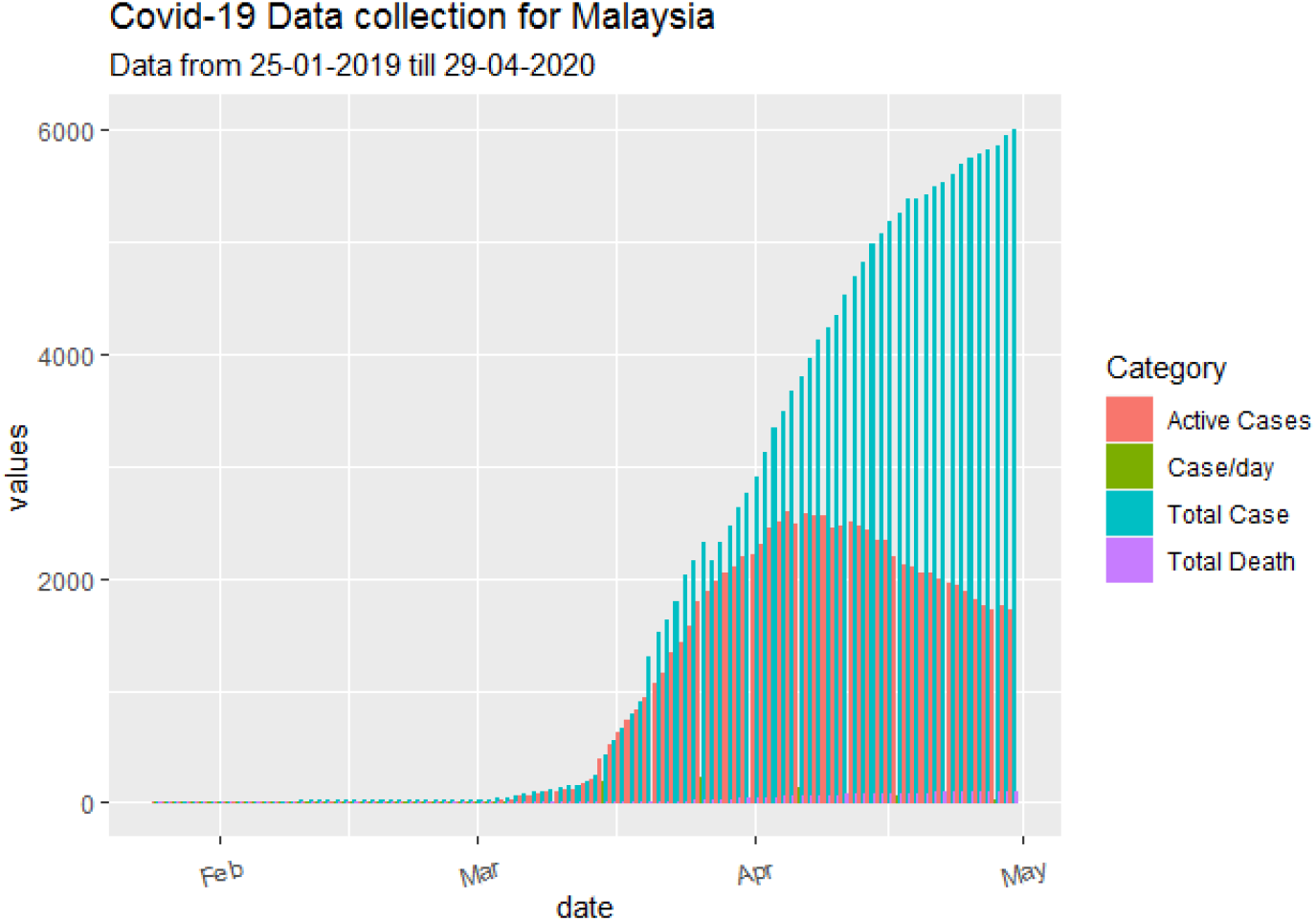
Total number of cases (including Total active cases, total death, total cases per day) for Malaysia.

Another visualization was performed based on state-wise cases which were used to understand the pattern of Covid-19 spread. Also, in Malaysia cases were divided as per based on clusters. That means each cluster is the major source of spreading the COVID-19 virus.

As shown in Figure 3(a), it is a visualization of all confirmed cases which were reported in January. As per sources, the first case was reported on 25th January 2020 in Malaysia. The first 3 cases were reported in Selangor state and 1 in Johor, later followed by Kedah. Figure 3(b) shows the total death for January. Two states were first to report first death cases in Malaysia, those are Johor and Selangor.

**Fig 3.**
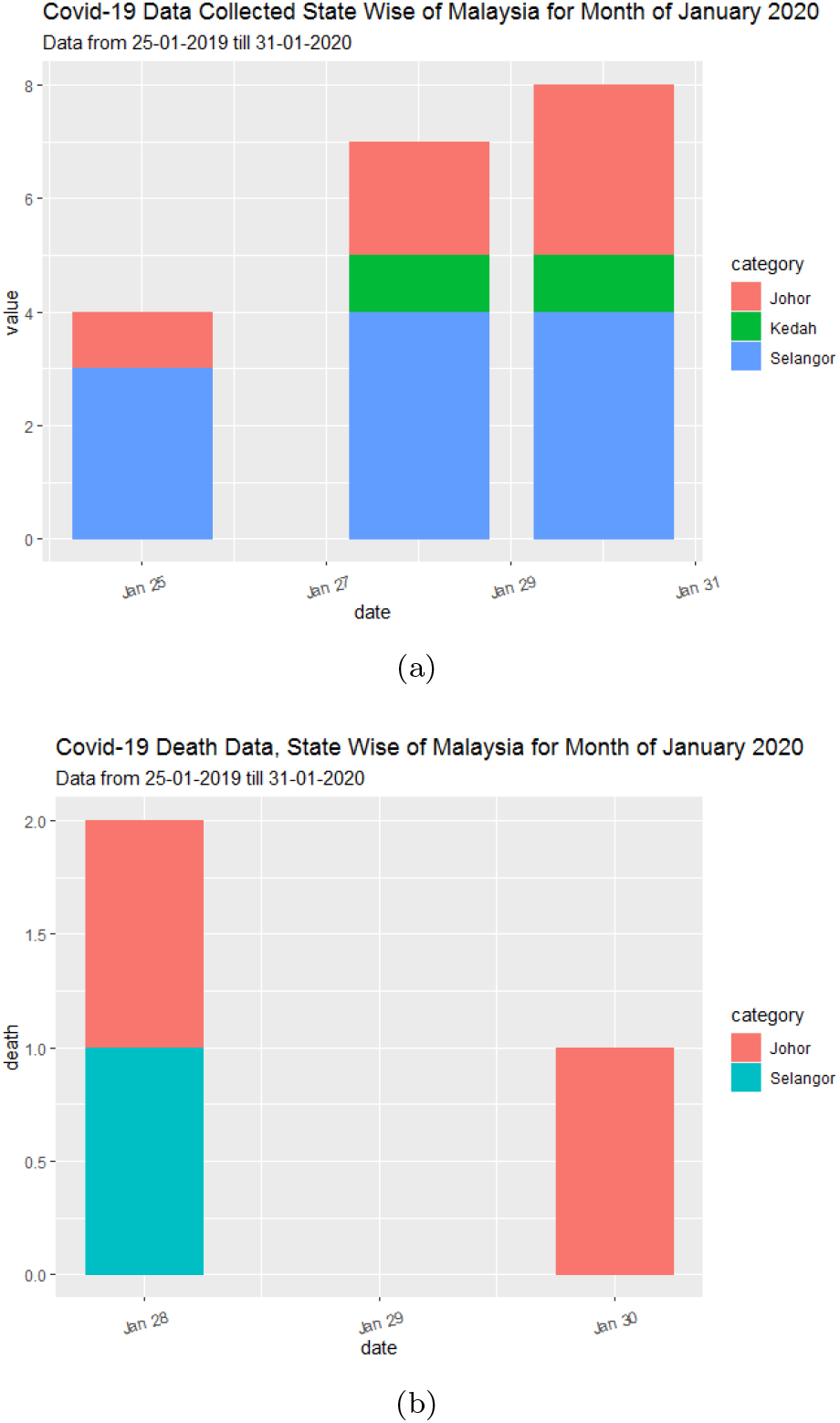
(a) State wise plotting and visualization of total confirmed cases for month of January 2020 (b) Total death cases reported for month of January 2020 bases on states of Malaysia.

Figure 4 the visualize the total cases reported for February 2020. Figure 4(a) displays all states where cases were reported based on clusters The cases were reported in Johor, Kedah, Kuala Lumpur Negri Sembilan, and Selangor. Among which Selangor and Kuala Lumpur played a major role in spreading the virus as shown in Figure 4(b). Indeed few deaths were counted for February which is plotted in Figure 4(c). Now looking at the cluster which was responsible to spread major outbreak in Malaysia. As per records from Feb 27 to March 3 14,500 Malaysian and 1,500 others attended a tabligh event “religious event” which were held at at Masjid Sri Petaling, Kuala Lumpur. On 9th March, Brunei’s first case was confirmed who traveled back from this event.

**Fig 4.**
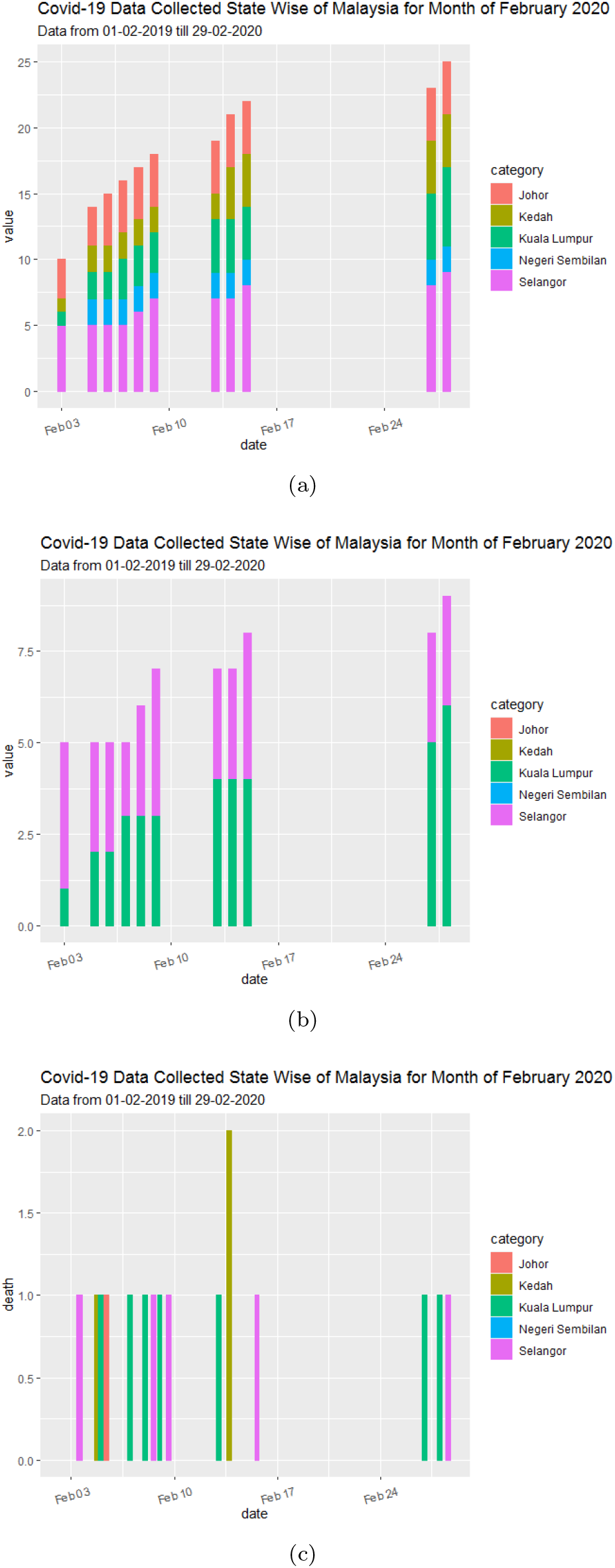
(a) State wise plotting and visualization of total confirmed cases for month of February 2020 (b) Maximum number of cases reported state wise for February 2020 (c) Total death cases reported for month of February 2020 bases on states of Malaysia.

Since at this stage it was a very big challenge to do testing of all participants and relatives of participants who attended this event. After the first cases were reported, the Malaysian government took a very important and game-changing step to enforce Movement Control Operation aka MCO. MCO was implemented in March 2020, to ensure the movement of all infected participants to restrict so that spread can be control and also social distance was enforced.

As shown in Figure 5 it represents the total number of confirmed cases based on state and number of death reported. In Figure 5(a) Total confirmed cases are visualized based on state-wise reporting, in this month all the 16 states were reported with virus spread. Whereas the maximum cases were reported from Kuala Lumpur, Selangor, Putrajaya. The Selangor cases were tacked back from the wedding which took place in Bangi Selangor on March 6-7. It was reported this cluster was the second in recording the number of cases after tabligh event. In this cluster, around 92 cases were reported, from which 44 cases were reported from Hospital Teluk Intan. Even though MOH believes, tabligh cluster is also linked with this cluster in spreading virus unknowingly.

**Fig 5.**
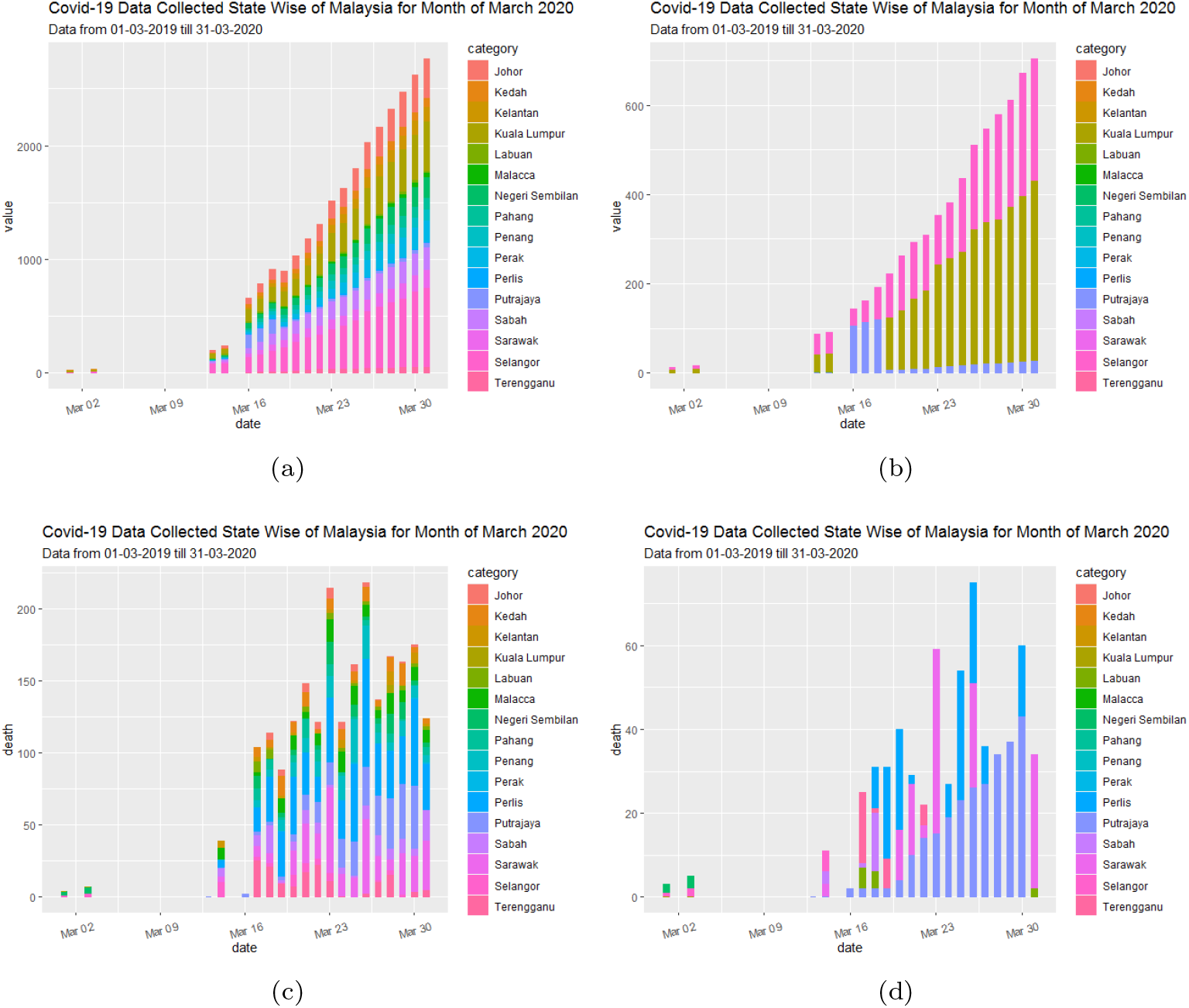
(a) State wise plotting and visualization of total confirmed cases for month of March 2020. (b) Maximum number of cases reported state wise for March 2020. (c) Total death cases reported for month of March 2020 bases on states of Malaysia. (d)Maximum number of death cases reported state wise for March 2020.

In the month of April maximum number of confirmed cases were reported. As shown in Figure 6(a) upto 6000 cases were notified. Figure 6(b) shows the maximum number reported based on states, Kuala Lumpur had reported maximum number of cases followed by same trend from last month, i,e. Selangore and Putraja. Also all reported death are plotted in Figure 6(c). Here a pattern can be mapped by looking at maximum number of death based on state which are Kuala Lumpur, Selangor, Sarawak, Negri Sembilan. These all states were infected badly by virus spread. Since, in this month MCO was on it ts third stage.

**Fig 6.**
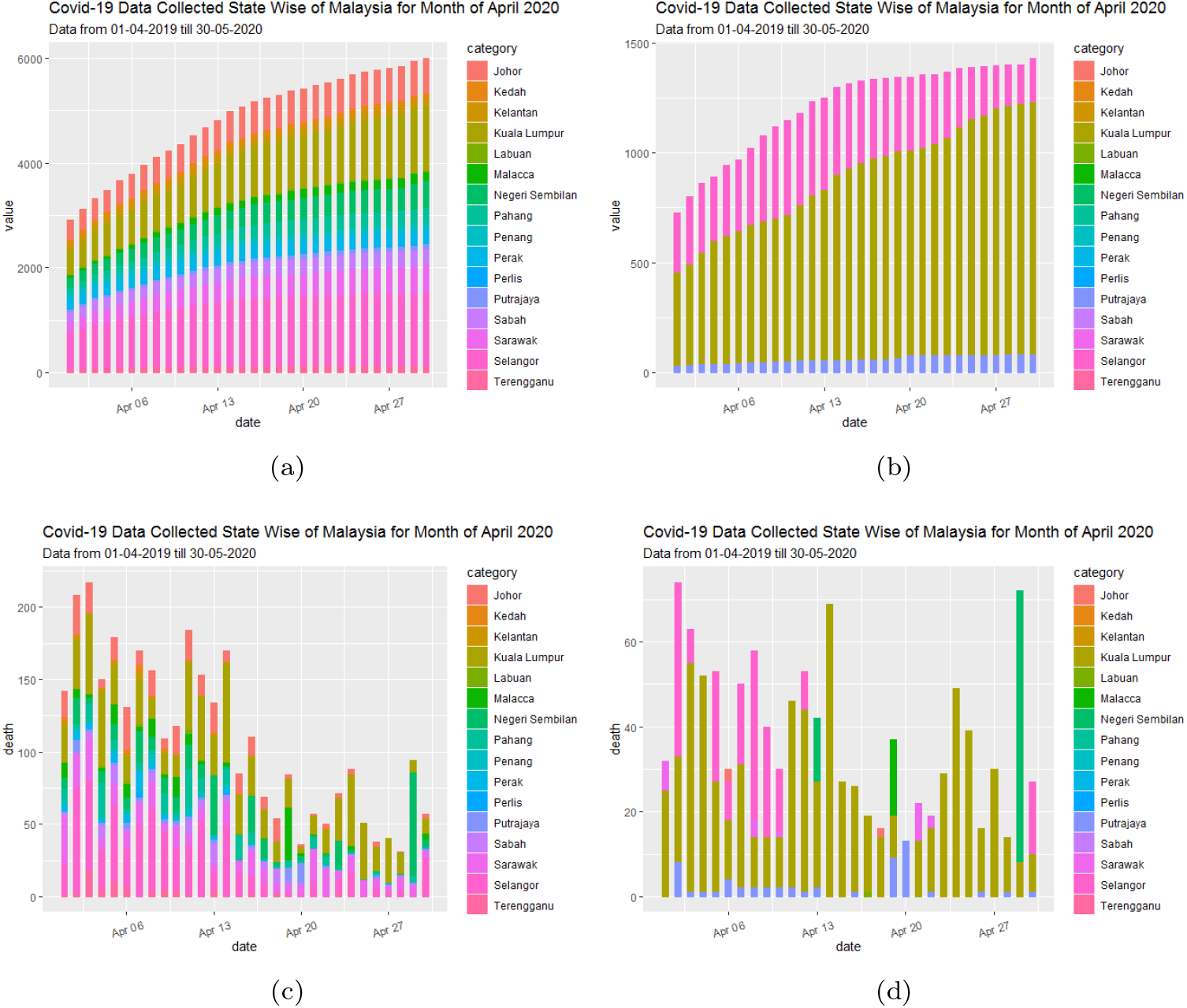
(a) State wise plotting and visualization of total confirmed cases for month of April 2020. (b) Maximum number of cases reported state wise for April 2020. (c) Total death cases reported for month of April 2020 bases on states of Malaysia. (d)Maximum number of death cases reported state wise for April 2020.

In April and May the cases reporting was controlled, as there were not bundle of cases were confirmed. As this can be linked with the MCO, hence it is very true to say only by applying social distancing and performing MCO can only provide risk-free Malaysia. Since in many states and districts there were no new cases recorded after the fourth stage of MCO applied. In Figure 7 (b) Only cases can be mapped in Kuala Lumpur, Selangor with limited cases, and Putrajaya with least reported cases for May 2020. Since the death rate also decreased as shown in Figure 7(c), also Figure 7(d) shows the major state where death cases were reported.

**Fig 7.**
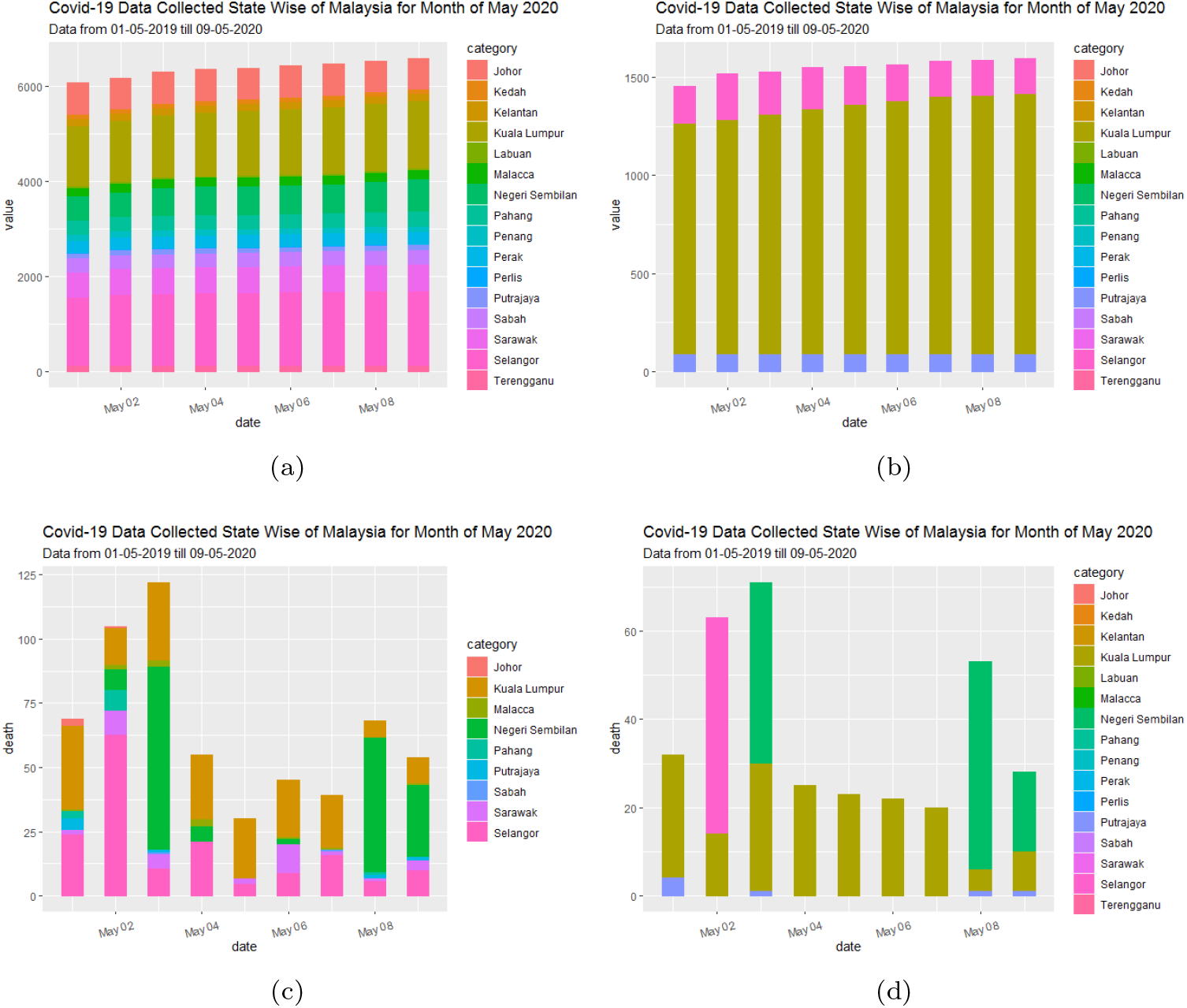
(a) State wise plotting and visualization of total confirmed cases for month of May 2020. (b) Maximum number of cases reported state wise for May 2020. (c) Total death cases reported for month of May 2020 bases on states of Malaysia. (d)Maximum number of death cases reported state wise for May 2020.

Since it is required to understand the pattern of virus spread as there is no clinically proven medication for this virus to cure. Hence, it is required to follow the social distancing and actively practice MCO until June 2020. In this paper, the next section will define how mathematically prediction can be developed to stay safe and how long MCO should be practiced.

## Methodology

The main objective of this paper is to predict the course of the spread of the COVID-19 pandemic in Malaysia. The paper uses the well-known SEIR model for estimating the number of the active cases on the time period between 10 March to 8 September 2020. Given the prevention measures imposed by the Malaysian Government, the paper investigates two scenarios, the first scenario attempts to estimate the number of active cases while considering enforcement of insufficient prevention measures. This scenario is implemented using parallel *Rt* values for the predefined time intervals. The second scenario attempts to estimate the number of active cases while considering strict enforcement of sufficient prevention measures. To mimic the effects of the imposed control measures, this scenario is implemented using sequential *Rt* values.

### Assumptions and Implementation

The main assumption in this manuscript is no local cases infected from animal transmissions, hence the initial cluster and the clusters that follow are from overseas. Moreover, we assume that the immunity of the population is the same. Finally, we assume that the natural death and birth have minimal impact on the course of predication, thus, they are not considered in the calculation.

In this paper, we also assume that the reproduction number Rt follows the strictness of the prevention measures imposed by the government. Thus, the extension of the lock-down decreases the value of the *Rt* over time. Accordingly, the course of the estimation continues with two assumptions; the first is the extension of the lock-down beyond 30 June 2020, and the second assumption is the gradual lift of the lockdown after 30 June 2020. Given the two assumptions above, the results attempt to estimate the number of active cases in Malaysia.

### The SEIR Estimation Model

The reason behind using a mathematical modeling of infectious disease is to help inform public health interventions, predict the future course of an outbreak and more importantly, to evaluate the strategies to control an epidemic in order to promote evidence-based decisions and policy [21]. According to the theory of compartmental models, the population is assigned to compartments with labels (i.e. S; refers to susceptible, E; refers to exposed, I; refers to infectious, and R; refers resistant or removed). The candidates of the population may progress between these compartments based on the characteristics of the infectious disease. In that context, the SEIR model belongs to the compartmental models’ family, which used to project how infectious diseases progress and show the likely outcome of an epidemic [39]. SEIR model is utilized successfully in estimating the number of infectious cases with pandemics such as Ebola [22] and SARS [23].

According to SEIR model, the population is classified into four compartments: susceptible (S; at risk of contracting the disease), exposed (E; infected but not yet infectious), infectious (I; capable of transmitting the disease), and removed (R; those who recover or die from the disease). Then, the total population (N) is given by the sum of the various compartments; *N* = *S* + *E* + *I* + *R*. The susceptible individuals who have been infected first enter a incubation period (exposed), where during this period, they are not likely to infect other individuals. The differential equations of the SEIR model are given as [42, 43].

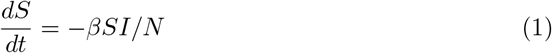

The transmission rate (*β*) is given by:

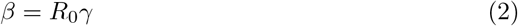

While the estimates of the infectious, exposed, and removed portions of the population are given by the following differential equations [42, 43]:

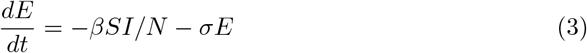

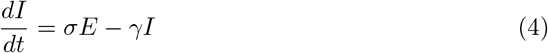

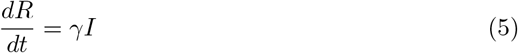

where *σ* is the infection rate calculated by the inverse of the mean incubation period, and *γ* is the recovery rate calculated by the inverse of infectious period.

The estimation is carried out for Malaysia with the major outbreak on 10 March 2020, where the number of infectious reached 129. According to the typical SEIR model, S was assumed to be the population of Malaysia. The initial number of cases as well as the respective recovered cases on 10 March 2020 are also considered. The number of exposed individuals is assumed to be 20 times the infectious individuals; *E* =20 * *I* [46]. According to [47], the infection rate (*σ*) was set as 1 = 5 : 2, where the denominator (5.2 days) is the average incubation period of COVID-19. The recovery rate (*γ*) is assumed to be 1/18, where the denominator (18) is calculated based on the summing the median time from infection to diagnosis plus the hospitalization period [48].

## Results and Discussion

In this paper, we follow the initial values of *Ro* published by [46] to estimate the number of active cases in Malaysia between 10^*th*^ March to 8^*th*^ September 2020. The first scenario assumes that the spread of the virus across the country with minimal prevention measures imposed. The second scenarios assumes the prevention measures are actively reducing the spread of the virus across the country. Thus, a sequential set of *Rt* are used to mimic the dynamics of the infection [46].

### The Impact of Imposing Minimal Prevention Measures

Despite the fact that this scenario is not realistic as active prevention measures are imposed all over Malaysia since 18 March 2020, the presented results provide insights for the policy makers on dynamics of the spread of the virus. Here, the active cases predication implements a parallel *Rt* values, where *R*_0_ is assumed to be 0.9, 2.6, and 3.1. Figure 8 shows that the number of the active cases will continue to climb for the whole time period, and the increase in the value of the reproduction number value *Rt* result in an exponential growth of the number of active cases. According to this scenario, the peak will occur by the end of July 2020, in case of an *Rt* = 3.1 which indicates not enforcing any form of prevention. By the end of August 2020, the number of active cases in Malaysia would be as follows 1012, 4600872, and 5979600, with respect to the chosen *Rt* values.

**Fig 8.**
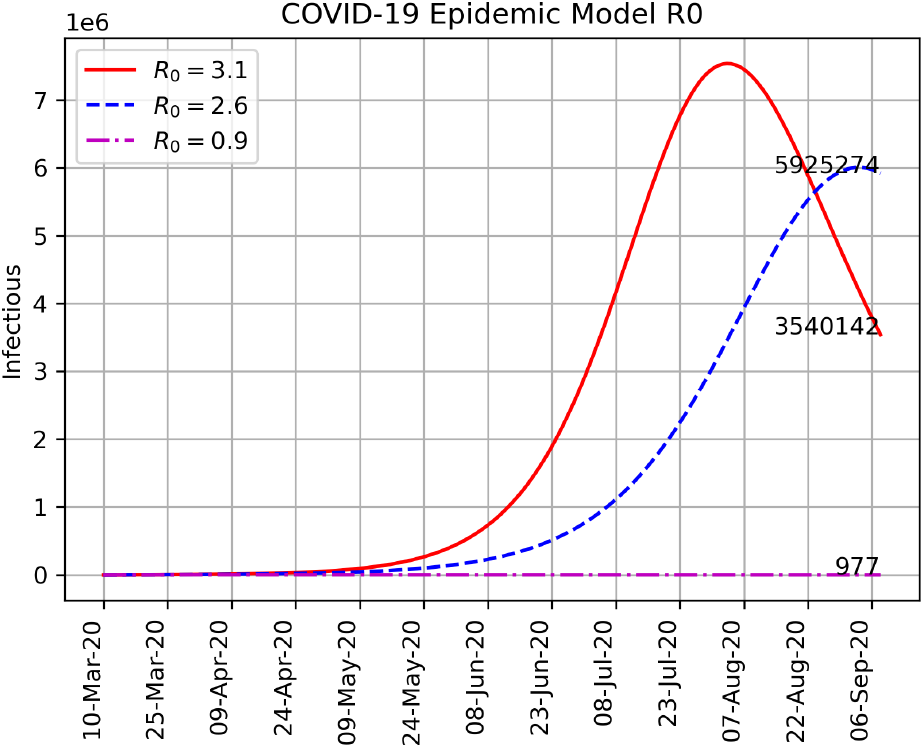
Three scenarios using parallel *Rt* values considering a minimal prevention measures.

### The Impact of Imposing Strict Prevention Measures

In order to consider the impact of the strict continuous lock-down with mass testing taking place on hot zones, we have dealt with two assumptions; the first is the extension of the lock-down beyond 10 June 2020, and the second assumption is the gradual lift of the lock-down after 10 June 2020. However, both assumptions follow the exact estimation patterns for the time period between 10 March to 10 September 2020.

The reason for choosing the 10 of March as starting date is due to the major outbreak occurred at that date, where the number of active cases spiked rapidly due to the discovery of multiple clusters across the country. Particularly, during the first stage (10-18 March 2020) a minimal control measures were implemented, followed by movement control order (MCO). However, the MCO is updated on 25 March, 10 April and 23 April receptively. The number of active cases in Malaysia reached the peak in mid-April as anticipated by the WHO. The reproductive number used to estimate the active cases during this period set as *Rt* = 2.9, 2.1, 0.9, 0.7, 0.5, 0.3, and 0.2 for stage 0, 1, 2, 3, 4, 5 and 6 respectively as shown in Figure 9.

**Fig 9.**
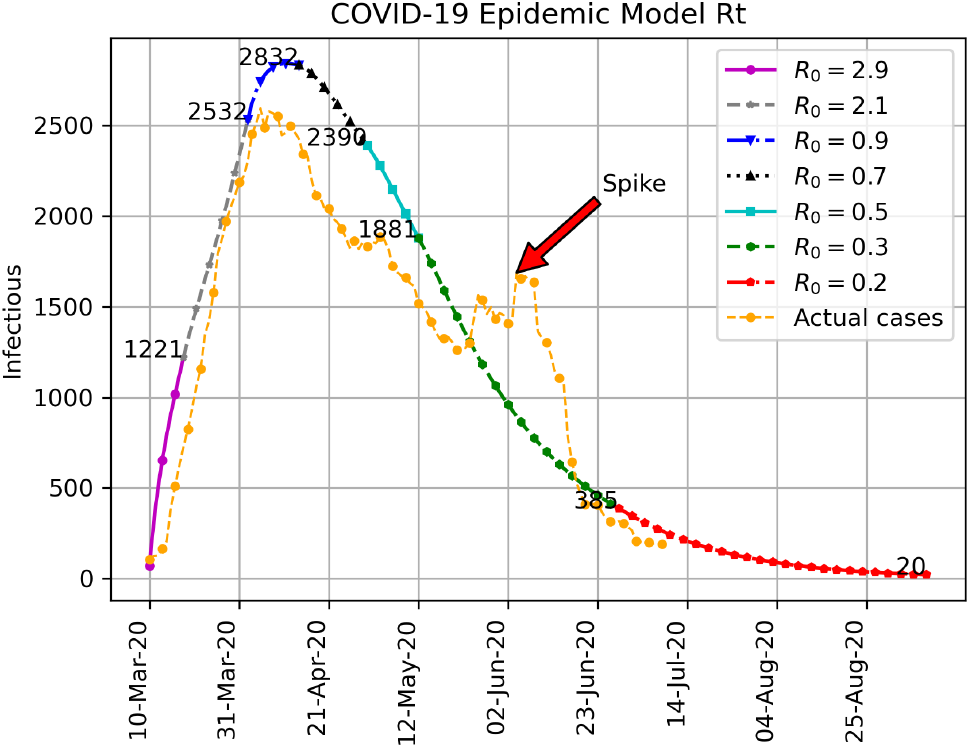
The prediction of active cases in Malaysia using sequential *Rt* values while considering the impact of the prevention measures.

The results shows that the Rt values is assumed to decrease gradually from 2.9 to 0.2 in Malaysia. The number of actual active cases reported on 28 June 2020 are 195, while the estimated active cases are 385. The variation between the reported and estimated active cases can be attributed to the infected cases showing mild symptoms or without showing symptoms which present no urgent need to have a medical check. The second scenario considers a gradual lift of the RMCO in the time span between (30 June to 31 August 2020). The second scenario considers a gradual lift of the RMCO in the time span between (30 June to 31 August 2020). Within this time period, three possibilities are considered based on the impact of the gradual lifting of the RMCO which result in a change in the value of the reproductive number applied in the estimation. Different reproductive numbers *Rt* = 0.7, 0.9, *and* 1.1 are used to mimic the impact of aforementioned gradual lifts of the RMCO during stage 6, see Figure 10(a), 10(b), and 10(c).

**Fig 10.**
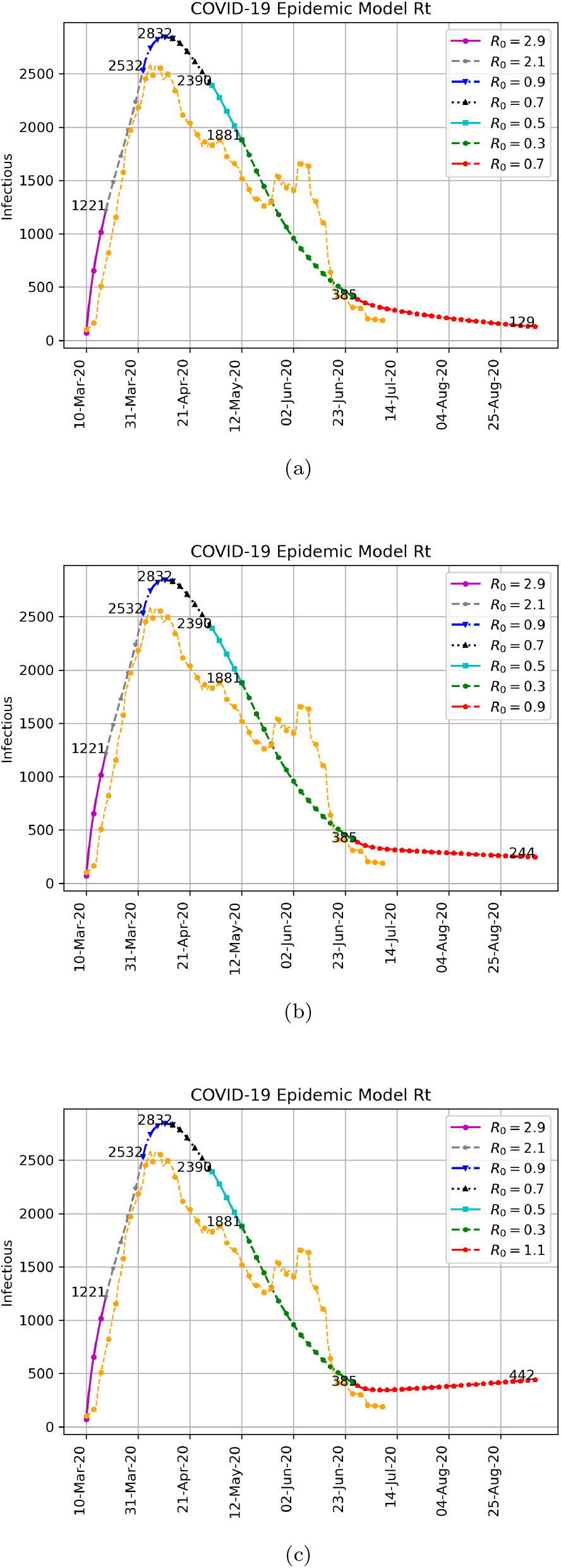
(a) The prediction of active cases in using *Rt* = 0.7 for the period between 10^*th*^ March – 30^*th*^ August 2020. (b) The prediction of active cases in Malaysia using *Rt* = 0.9 for the period between 10^*th*^ March – 30^*th*^ August 2020. (c) The prediction of active cases in Malaysia using *Rt* = 1.1 for the period between 10^*th*^ March – 30^*th*^ August 2020.

The number of estimated active cases on 31 August 2020 are 138 (Figure 10(a)), 1605 (Figure 10(b)), and 1817 (Figure 10(c)), with respect to *Rt* = 0.7, 0.9, and 1.1 respectively.

These estimated numbers of the active cases depend heavily on the prior estimate of the active cases on 12 May 2020. The variations among these estimates are attributed to the differences in the used *Rt*.

## Discussion

In this paper, different scenarios of the spread of Covid-19, are modeled to reflect the number of active cases in Malaysia. The applied SEIR model depends on the accuracy of the parameters used in the estimation, for instance, the incubation period and the reproductive number values.

Based on the outcome of the SEIR model and the actual data of active cases published by the Ministry of Health, Malaysia, we can easily see the impact of the sequential MCO on the course of the spread of the pandemic. In that context, the initial stage considered in this paper shows that the initial value of the reproduction number is set to 2.9. The estimate obtained by the SEIR model fits the actual data collected from the official portal of the Ministry of Health Malaysia, where the number of active cases by the end of this stage reaches 1221 and shows a clear exponential growth. Moreover, the first stage of the MCO (18 to 30 March) has reduced the *Rt* value from 2.9 to 2.1 which reflected in a reduction of the active cases to around 2000 patients.

The second stage of the MCO (30 March to 14 April) was capable to reduce the Rt value from 2.1 to 0.9. This show that the prediction matches the actual data and accordingly, the peak of the active cases curve occurred in mid-April (anticipated by the WHO). Similarly, the further extension of the MCO result in a noticeable decline in the active cases curve, where by 12 May 2020, the estimate of the active cases is 1881 as compared to 1410 obtained from the actual data.

The paper also further extend the prediction to estimate the number of the active cases till 8^*th*^ September 2020. In order to simulate the impact of the relaxation of the RMCO, the paper used a sequence of various *Rt* values. Table 2 summarizes the outcomes of the prediction. The table considers the *Rt* value with respect to the estimated number of active cases.

**Table 2.** A summary of the active cases prediction in Malaysia till 8 July 2020.

| $R_t$ Values | Estimate of the active cases by the end of September 2020 | Description |
| --- | --- | --- |
| 0.3 | 28 | Extension of strict MCO |
| 0.7 | 138 | Extension of a relaxed relaxed MCO |
| 0.9 | 238 |  |
| 1.1 | 395 |  |

The various *Rt* values matches the severity of the imposed RMCO, where keeping the RMCO in its current form will reduce the number of active cases to 28 case by the end of August 2020.

It is learned that Malaysia’s strategy to break the chain of infection appears to be successful with high number of cases (2532 case) during March 2020, a sudden decrease was found on the 15^*th*^ April itself, with only 85 cases. Besides, the percentage of recovery also was found to be high, exceeding 50%.

The predictions made by this paper were aligned with the health authorities’ source of information and public opinion. Thus, the extension of the EMCO, from 15^*th*^ April to 28^*th*^ April was very crucial and successfully prohibited the increase of COVID-19 cases, breaking the chain of infection and flattens the curve.

## Conclusions

The conducted study attempted to estimate the number of active cases of COVID-19 in Malaysia between 10 March to 8 September 2020. The study applies a sequential reproductive number values to mimic the dynamics of the virus spread given the strict prevention measures imposed by the Malaysian authorities. The limitations and future extension of the study can be summarized as follows; The value of the reproductive number (as well as the rest of the model’s parameters) might be computed adaptively given the available information such as the density of the population, the average contact time. The country-wise estimate might be predicted more accurately by acculturating the estimates of the province/states. Moreover, additional consideration should be given to the hot spot across the country given the current model lacks this feature. In the case of Malaysia, it was eagerly handled carefully to control the number of cases. Malaysia’s government did all possible ways to stop the spread of COVID-19. After the spike notified in March 2020, MCO was announced in whole Malaysia to control spread.

It was reported by the government after the control of cases for till April 2020, another spike was notified in the month of April 2020 (last week) and June 2020(early weeks), in which most of the cases were reported from immigrants as shown in Figure 9. These all immigrants were kept in the detention center. This may conclude that COVID-19 is not spread among Malaysian as these immigrants were all living without following social distancing parameters.

## Acknowledgments

Author wants thanks Universiti Teknologi PETRONAS for providing resources. Author also want to show gratitude to all co-authors for providing their support and help in conducting this research.

This research was funded by Yayasan Universiti TEknologi PETRONAS grant number (YUTP 015LC0-299)”

